# DIFFERENTIAL BIOENERGETICS IN ADULT RAT CARDIOMYOCYTES ISOLATED FROM THE RIGHT VERSUS LEFT VENTRICLE

**DOI:** 10.1101/2020.06.08.133769

**Authors:** Quyen L. Nguyen, Krithika Rao, Steven J. Mullett, Stacy G. Wendell, Claudette St. Croix, Eric Goetzman, Sruti Shiva

## Abstract

The right and left ventricle of the heart have distinctly different developmental origins and are affected differently by similar pathological stimuli. Though it is well established that the heart relies almost entirely on mitochondrial function to sustain energy production, it remains unclear whether bioenergetics differ in the two ventricles. Herein, we define a novel methodology to optimize the isolation of intact cardiomyocytes from the right versus the left ventricle. We demonstrate that this segmental Langendorff-free methodology yields viable cardiomyocytes with intact mitochondrial function. Further, we compare bioenergetics in right versus left ventricle cardiomyocytes and show that cardiomyocytes from the right ventricle have a greater maximal capacity for respiration and enhanced glycolytic rate. This increase in respiration was concomitant with increased fatty acid oxidation and levels of fatty acid oxidation proteins, but no change in mitochondrial electron transport complex expression. These data validate a potentially powerful tool to evaluate differences in right and left ventricular function and advance the understanding of cardiac bioenergetic differences. These data will be discussed in the context of differential responses by the right versus ventricle in pathology.

## 1. INTRODUCTION

Right ventricular (RV) failure is a terminal consequence of cardiopulmonary disease, including pulmonary vascular disease, lung disease, and left heart disease. The function of the RV is a major determinant of morbidity and mortality in these conditions.^1-6^ Accumulating evidence suggests that the mechanisms that underlie RV failure may diverge from those that occur in failure of the left ventricle (LV), thus underscoring the need for dedicated study of the RV.^7-10^

Experimentally, the RV tolerates acute pressure overload more poorly compared to the LV.^11^ Clinically, the time course to RV failure appears contracted compared to LV failure. Patients survive for ∼14 years with systemic hypertension and progressive LV hypertrophy before the onset of apparent congestive heart failure symptoms.^12^ In contrast, upon the diagnosis of pulmonary hypertension ∼4 years after symptom onset, 94% of patients already have apparent RV failure symptoms.^13^ Mechanistic differences between the RV and LV response to stress remain unclear, and extrapolation of LV biology towards the RV may be limited by inherent ventricular differences in origin, structure, and function.^9,10^

As the most metabolically active organ with the highest cellular mitochondrial density, the heart continuously produces adenosine triphosphate (ATP), ∼90% of which is derived from mitochondrial oxidative phosphorylation.^14^ The mitochondrial dysfunction that has been correlated to the bioenergetic collapse seen in heart failure derives largely from studies in LV disease,^15^ and is characterized by depressed oxidative phosphorylation,^16,17^ a metabolic switch from fatty acid oxidation (FAO) to glycolysis,^18^ lipid accumulation and lipotoxicity,^19^ and changes in mitochondrial morphology and dynamics.^20-22^ Some of these changes have been recapitulated in models of RV failure, particularly the switch from FAO to glycolysis.^23-25^ Beyond this, few studies directly compare the metabolic differences between the LV and RV in health and disease. In dogs, the LV was shown to respond to increased myocardial work by augmenting coronary blood flow, while the RV was shown to augment blood flow and oxygen extraction.^26,27^ Studies elucidating the molecular basis for this ventricular metabolic difference are sparse. Genome-wide transcription profiling in mouse^28^ and rat^29^ reveal LV versus RV differences in gene expression, including those genes involved in metabolism of carbohydrates, lipids, and nucleic acids.^29^ Bruns et al showed in control calves increased RV versus LV mitochondrial content and expression of COXI (mitochondrial complex IV) and Mfn1, which regulates mitochondrial fusion.^7,8^ Taken together, these studies suggest a mitochondrial basis for ventricular differences in metabolism.

The study of cardiac mitochondrial function in relevant biological models requires viable primary cells, which are historically challenging to procure and maintain in culture. Isolation of primary adult cardiomyocytes utilizing the Langendorff apparatus demands significant investment in specialized equipment and training, as well as lengthy experiment times. Building upon a Langendorff-free approach for cardiomyocyte isolation recently described by Ackers-Johnson et al. in the adult mouse heart,^30^ we developed a methodology to segmentally isolate primary cardiomyocytes from separated adult rat LV and RV. Utilizing cells obtained by this technique to directly compare cellular metabolism between ventricles, we show distinct bioenergetic profiles between the LV and RV myocytes, which are accompanied by differences in substrate metabolism and mitochondrial enzymatic function.

## 2. METHODS

### 2.1 Isolation of adult rat cardiomyocytes

The experimental protocol was in accordance with the Guide for the Care and Use of Laboratory Animals and approved by the University of Pittsburgh IACUC (Protocol #15127354). Male Sprague Dawley rats aged 9-12 weeks were used in all experimental procedures.

#### 2.1.1 Equipment

CO_2_ chamber

Heated stir plate

10 cm culture dishes (non-sterile)

Dissection scissors

Dissection microforceps

Straight-locking hemostat surgical clamps (for aorta clamping)

25-gauge hypodermic needles

20 ml syringes

250 µm tissue strainers (for 15 ml conical tubes)

Hemocytometer

#### 2.1.2 Protocol steps

1. Prepare calcium-free isolation buffer and collagenase digestion buffer as described in **Table 1**.
2. Anesthetize animals by CO_2_ at 1.11 L/min for 10 minutes, until cessation of respiration.
3. Using dissection instruments, open the chest to expose the heart and great vessels.
4. Using a 25-gauge needle, inject 15 ml of ice-cold calcium-free buffer into the RV to flush free of blood.
5. Identify and clamp the ascending aorta with a small straight-locking hemostat. Inject 0.1 ml of heparin (1000U/ml) into the LV apex to prevent coronary thrombosis.
6. Excise the heart from the chest cavity and inject 30-50 ml calcium-free buffer into the LV apex until the coronary arteries and effluent fluid (out of the left atrium) are clear of blood.
7. With the aorta still clamped, the heart can be placed into ice-cold calcium free buffer during the time needed to procure hearts from additional animals. In our experiments, up to four hearts were processed during a single session.
8. When ready for digestion, transfer hearts into a 10 cm dish atop a 37°C warmer containing 5 ml of collagenase buffer.
9. Inject collagenase buffer into the LV apex of each heart. We used ∼50 ml volume of buffer per two hearts, which was recirculated until digestion was achieved. We gauged progress of digestion by the softening texture of the heart, loss of pressure during collagenase injection, and presence of cells actively coming into solution during injection. The presence of myocytes can be confirmed by microscopic examination of a small volume of fluid during the digestion process.
10. After digestion, remove the atria and separate the RV free wall from the LV with scissors. For each heart, place the two ventricles into separate dishes.
11. Wash the tissue in 0.5 – 1 ml ice-cold calcium free buffer. We did not find that the addition of fetal bovine serum (to stop digestion) affected cell yields.
12. Tease apart the digested tissues with microforceps into 1 mm pieces and triturate gently with a pipette to further dissociate cells into solution. Pipette tips were cut to produce a larger opening to minimize shear stress to cells during trituration.
13. Pass LV and RV myocyte suspensions through separate 250 µm meshes into 15 ml conical tubes and allow cells to settle by gravity. Viable myocytes preferentially settle by gravity first, while non-viable myocytes and non-myocyte cells (e.g. fibroblasts) remain in the supernatant.
14. Reintroduce calcium by four sequential additions of Dulbecco’s Modified Eagle Medium (DMEM, containing 1.8 mM CaCl_2_), until the final volume is doubled. At this point, the suspension will be composed of 1:1 calcium free buffer and DMEM with a calcium concentration of 0.9 mM. Allow for gravity settling of cells in between each addition of DMEM to produce a myocyte-rich fraction within the pellet.
15. Remove excess supernatant to concentrate the cell suspension.
16. Count cardiomyocytes by a hemocytometer.
17. For extracellular flux analysis, cells are plated at 15,000/well density onto a laminin-coated 96-well plate and allowed to adhere. After about 1-2 hours of attachment, change the media to bicarbonate-free DMEM (containing 1.8 mM calcium).
18. Cells not utilized for same-day experiments can be frozen at −80°C for later analyses.

**Table 1:** Composition of buffers for cardiomyocyte isolation.

| **Calcium-free isolation buffer** | Molecular Weight<br>(g/mol) | Concentration<br>(mM) | g/500 mL MQ H<sub>2</sub>O |
| --- | --- | --- | --- |
| NaCl | 58.44 | 130 | 3.8 |
| KCl | 74.55 | 5 | 0.186 |
| NaH<sub>2</sub>PO<sub>4</sub> | 119.96 | 0.5 | 0.03 |
| HEPES | 238.3 | 10 | 1.19 |
| Glucose | 180.16 | 10 | 0.91 |
| BDM | 101.11 | 10 | 0.506 |
| Taurine | 125.15 | 10 | 0.626 |
| MgCl<sub>2</sub> | 95.2 | 1 | 0.048 |
| **Collagenase buffer** | Activity units*<br>(IU/mg) | Target Activity/ml | mg/50 mL<br>calcium-free<br>isolation buffer |
| Type 2 collagenase<br>(Worthington Biochemical<br>Corporation, Lakewood, NJ) | 286 | 300 | 45 |

\*activity of collagenase varies from batch to batch. Amount of collagenase used is determined by desired activity units/ml

### 2.2 Live cell imaging and mitochondrial staining

Differential interference contrast (DIC) images of live cardiomyocytes from separated LV and RV were captured at 60X by a Nikon Eclipse Ti2 microscope (Nikon Instruments, Melville, NY). Tetramethylrhodamine, methyl ester (TMRE; 100 nM) was used to highlight active mitochondria with intact membrane potentials within myocytes. The mitochondrial membrane potential was depolarized using carbonyl cyanide p-(trifluoro-methoxy) phenyl-hydrazone (FCCP; 1 µM).

### 2.3 Quantification of TMRE fluorescence

Cell suspensions of LV and RV myocytes were incubated with TMRE (100 nM) and plated at 10,000 and 20,000/well density into a 96-well plate. TMRE fluorescence was quantified before and after addition of FCCP spectrophotometrically using a Synergy H1 plate reader (Biotek) with the absorbance peak at 548 nm and emission peak at 574 nm.

### 2.4 Measurement of cardiomyocyte bioenergetics

Oxygen consumption rate (OCR) and extracellular acidification rate (ECAR) were measured in isolated cardiomyocytes by extracellular flux analysis (XF96, Seahorse Biosciences, Billerica, MA). Typical cell yields by our isolation procedure were ∼1.5 × 10^6^ cells for each LV and ∼0.7 × 10^6^ cells for each RV, with 60-80% rod-shaped cells. Cardiomyocytes were plated at 15,000 cell/well density onto 96-well plates previously coated with laminin (60 ug/well) and allowed to attach for 1-2 hours, followed by change of media to bicarbonate-free DMEM for XF analysis. Optimal cell density for XF analysis was based on the protocol by Readnower et al.^31^ Cardiomyocytes further rested for an additional 1-2 hours prior to XF analysis.

After measurement of basal OCR, OCR due to proton leak was determined by oligomycin A (2.5 µM) treatment. Maximal uncoupled OCR was measured by the addition of the uncoupler carbonyl cyanide p-(trifluoro-methoxy) phenyl-hydrazone (FCCP; 1 µM). Non-mitochondrial OCR (defined as the oxygen consumption rate of all cellular processes excluding mitochondrial respiration) was measured in the presence of rotenone (10 µM). In a subset of samples, etomoxir (400 µM), UK5099 (2 µM), and Bis-2-(5-phenylacetamido-1,3,4-thiadiazol-2-yl)ethyl sulfide (BPTES; 3 µM) were added to quantify the OCR dependent on oxidation of fatty acid, glucose, and glutaminolysis, respectively. In a subset of samples, BSA-palmitate (100 µM) and carnitine (100 µM) were added to quantify the rate of exogenous fatty acid oxidation (FAO.) Glycolytic rate was calculated by determining extracellular acidification rate (ECAR) that was sensitive to 2DG. Following XF analysis, cells within individual wells were lysed for protein quantification.

### 2.5 Measurement of cardiomyocyte fatty acid oxidation

Fatty acid oxidation was measured using freshly isolated cardiomyocytes incubated in suspension with ^14^C-palmitate (Perkin Elmer) conjugated to BSA. Reactions contained 50,000 cells, 125 µM ^14^C-labeled palmitate, and 100 µM carnitine in a 200 µl total volume of serum-free DMEM. After 1 hr rotating at 100 rpm in a 37°C water bath, the reactions were stopped by addition of perchloric acid to a final concentration of 0.5 mM. The ^14^CO_2_ released by acidification was captured on filter papers soaked in 2M KOH and subjected to scintillation counting. Each LV and RV cell isolate was measured at least in duplicate.

### 2.6 Metabolomics by untargeted liquid chromatography high-resolution mass spectrometry (LC-HRMS)

Metabolic quenching and polar metabolite pool extraction was performed using ice cold 80% methanol/0.1% formic acid at a ratio of 500 µL per 1 × 10^6^ adherent cells. A deuterated internal standard mix that included (d_3_)-creatinine, (d_3_)-alanine, (d_4_)-taurine and (d_3_)-lactate (Sigma-Aldrich) was added to the sample lysates at a final concentration of 10 µM. After 3 minutes of vortexing, the supernatant was cleared of protein by centrifugation at 16,000 x g. Cleared supernatant (3µL) was subjected to online LC-HRMS analysis. Briefly, samples were injected via a Thermo Vanquish ultra-high performance liquid chromatograph (UHPLC) autosampler onto a Thermo HyperCarb porous graphite column (2.1×100mm, 3μm particle size) maintained at 55°C. For the 20 minute LC gradient, the mobile phase consisted of the following: solvent A (water/0.1% FA) and solvent B (ACN/0.1% FA). The gradient was the following: 0-1min 1%B, increasing to 15%B over 5 minutes, and to 98%B over 5 minutes. The gradient is held at 98%B for 5 minutes before equilibration to initial conditions. The Thermo ID-X tribrid mass spectrometer was operated in both positive and negative ion mode, scanning in Full MS mode (2 μscans) from 100 to 800 *m/z* at 70,000 resolution with an AGC target of 2 × 10^5^. Source ionization setting was +3.0 kV or −2.7kV spray voltage, respectively for positive and negative mode. Source gas parameters were 35 sheath gas, 12 auxiliary gas at 320°C, and 8 sweep gas. Calibration was performed prior to analysis using the Pierce™ FlexMix Ion Calibration Solutions (Thermo Fisher Scientific). For specific analytes, integrated peak areas were then extracted manually using Quan Browser (Thermo Fisher Xcalibur ver. 2.7). Untargeted analysis was performed by using Compound Discoverer (Thermo Fisher ver. 3.1) to align and quantify all peaks, defined as a unique mass to charge ratio at a unique retention time, from both polarities and search the results against database entries (mzVault, mzCloud, ChemSpider, and internal mass databases) for putative identifications. All analyte peak areas were then normalized to internal standard and cell number before being filtered by occupancy rate, allowing one missing value from a group in comparison. Next, statistics were applied via two-tailed t-test to compare groups with fold changes calculated from the mean of groups. Finally, *p* values and fold changes were log converted to generate volcano plots where analytes are ranked by significance.

### 2.7 Mitochondrial DNA

Total DNA was isolated from LV and RV cardiomyocytes and mitochondrial DNA copy number was quantified by RT-PCR using the primer for mitochondrial encoded gene ND1 (R: 5’ GGC GTC TGC AAA TGG TTG TAA; F: 5’ AAT CGC CAT AGC CTT CCT AAC AT). MtDNA was expressed as a ratio of nuclear DNA (nDNA) quantified by RT-PCR using the primer for 18S (F: 5” TTG ATT AAG TCC CTG CCC TTT GT’; R:5’ CGA TCC GAG GGC CTA ACT A) in cardiomyocytes.

### 2.8 Mitochondrial enzyme expression

Mitochondrial protein expression was measured by Western blot. Antibodies for complexes (Complex I #MS112, Complex II #MS204, Complex IV #MS407) were purchased from MitoSciences, now Abcam, Cambridge, MA. Antibodies for carnitine palmitoyltransferase-1A (CPT1A, ab128568), Complex V (ATP5A, ab14748), Mfn1 (ab126575) and β-actin (ab8226) were purchased from Abcam, Cambridge, MA. CPT1B (NBP1-59576) antibody was purchased from Novus Biologicals, Centennial, CO. Antibodies for VLCAD and LCAD were a kind gift of Dr. Jerry Vockley.

### 2.9 Mitochondrial enzyme activities

Following several cycles of freeze/thaw, CPT1 activity of isolated myocytes was determined by spectrophotometrically monitoring the generation of CoA-SH from 100 µM palmitoyl-CoA in the presence of 5 mM L-carnitine and 200 µM 5,5’-dinitro-bis-(2-nitrobenzoic acid) (DTNB) at an absorbance of 412 nm, as adapted from prior studies.^32^ Enzymatic activity of complexes I, II, IV, and citrate synthase was spectrophotometrically measured as previously described.^33^

### 2.10 Statistics

Statistics were performed on GraphPad Prism 8.0 (La Jolla, CA). Data in LV and RV from the same heart were compared by paired *t*-test or repeated measures analysis of variance (ANOVA) where appropriate. *P* < 0.05 was considered significant.

## 3. RESULTS

### 3.1 Myocytes isolated from separated LV and RV display intact mitochondrial membrane potential

Segmental isolation of cardiomyocytes yielded morphologically intact cells from the LV and RV, which have characteristic rod-shape and sarcomeric striations (**Figure 1A-B**) and TMRE fluorescence in these myocytes demonstrate viability and mitochondria with intact membrane potential which dissipates following addition of FCCP (**Figure 1B-D**).

**Figure 1.**
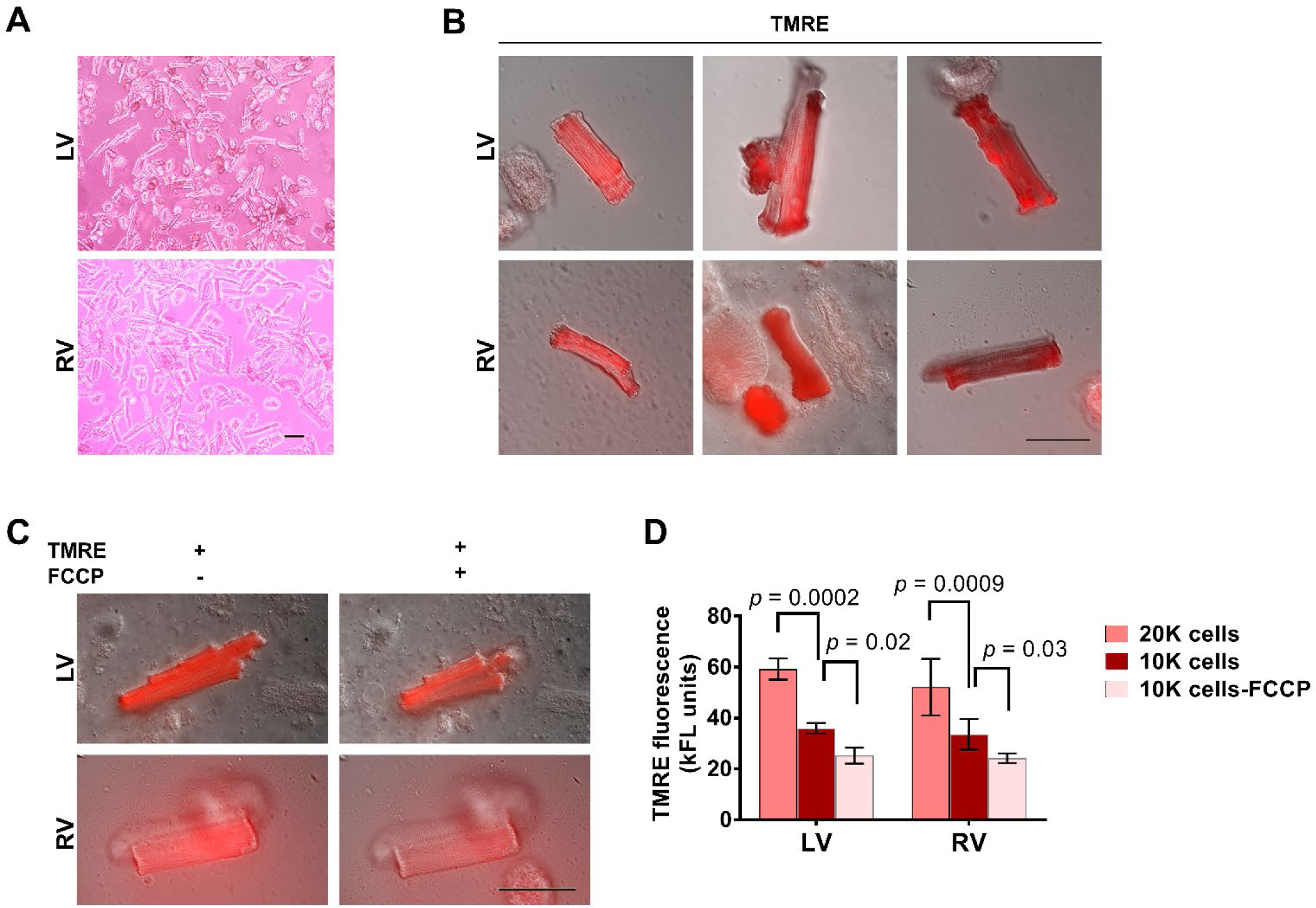
Isolation method yields viable cardiomyocytes from separated adult rat left and right ventricles. **A:** Representative light microscopy images at 10X of cardiomyocytes plated for extracellular flux analysis. Bar = 50 µm. **B:** Representative differential interference contrast (DIC) images at 60X of cardiomyocytes stained with TMRE from left and right ventricles. Bar = 50 µm. **C:** Dissipation of TMRE from representative cardiomyocytes following addition of the ionophore FCCP. **D:** Spectrophotometric plate reader quantification of TMRE fluorescence in cell suspensions of cardiomyocytes isolated from LV and RV from n = 5 animals at different cell densities and following addition of FCCP. Data are mean ± SEM. LV, left ventricle; RV, right ventricle; TMRE, Tetramethylrhodamine, methyl ester; FCCP, carbonyl cyanide p-(trifluoro-methoxy) phenyl-hydrazone.

### 3.2 Cardiomyocytes from LV and RV show distinct bioenergetic profiles

Extracellular flux (XF) analysis showed that both RV and LV cardiomyocytes exhibited appropriate response in oxygen consumption rate (OCR) to mitochondrial modulators (**Figure 2A**). After controlling for the non-mitochondrial (rotenone-insensitive) OCR, we found basal, ATP-linked, and proton leak OCR to be similar for myocytes from both ventricles. However, in the presence of the protonophore FCCP, RV myocytes showed significantly enhanced maximal uncoupled OCR compared to LV (5.3 ± 0.73 vs 4.0 ± 0.58 nmol O_2_/min/mg protein, *p* = 0.0002; **Figure 2B**). Accordingly, the reserve respiratory capacity (maximal OCR – basal OCR) was also significantly higher in RV myocytes (*p* = 0.032; **Figure 2B**).

**Figure 2.**
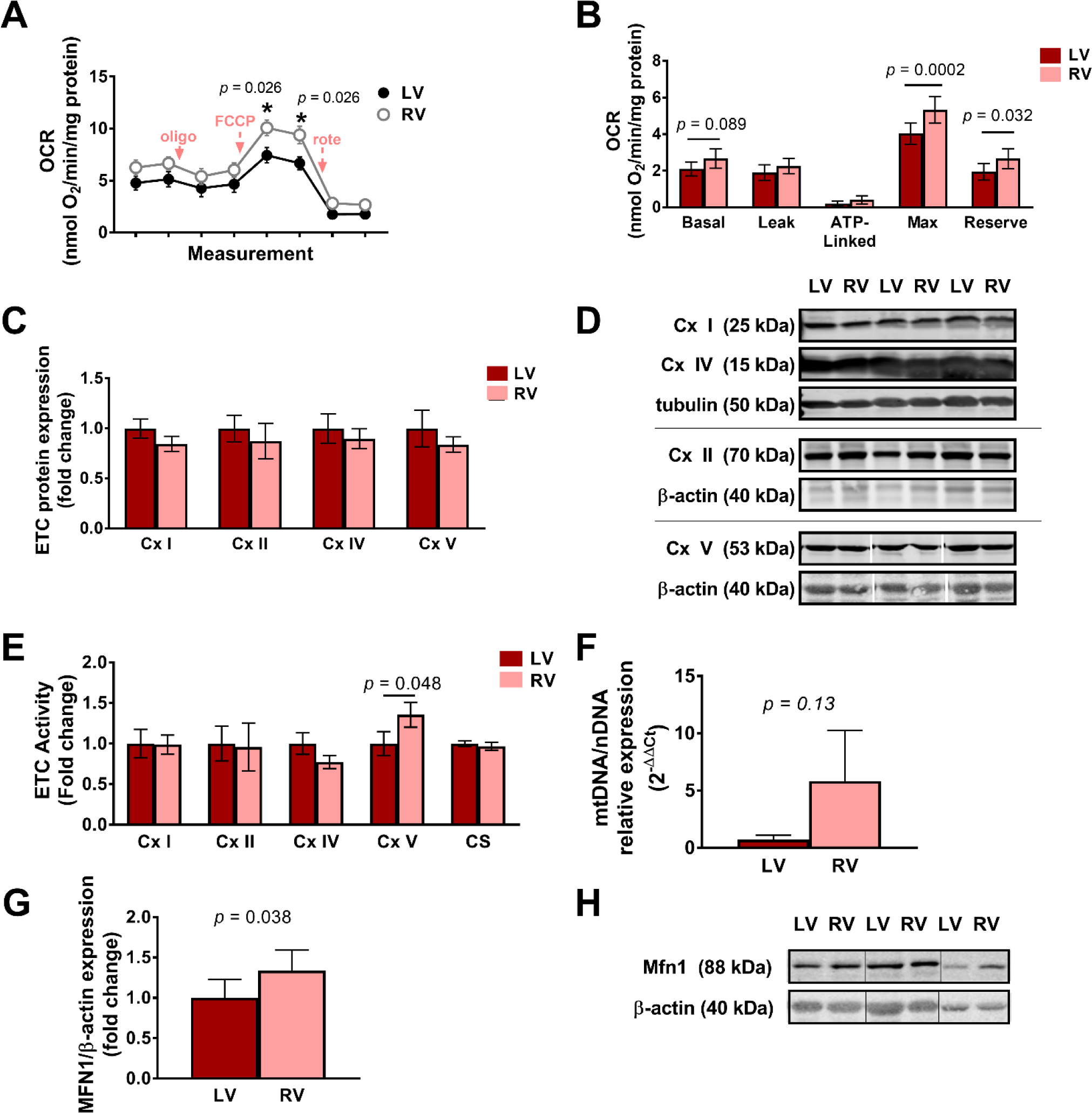
Separated LV and RV cardiomyocytes display distinct bioenergetic profiles. **A:** Oxygen consumption rate (OCR) of LV and RV myocytes measured in basal state, and following injections of oligomycin A (oligo), carbonyl cyanide p-(trifluoro-methoxy) phenyl-hydrazine (FCCP), and rotenone (rote). **B:** Individual components of the bioenergetic profile of LV and RV myocytes isolated from n = 20 animals. **C:** Pooled densitometric analysis and **D:** representative Western blots of protein expression of ETC complexes in LV and RV myocytes from n = 10-17 animals. **E:** Spectrophotometric measurement of kinetic activities of ETC complex (Cx) enzymes and citrate synthase (CS) in LV and RV myocytes from n = 6-17 animals. **F:** Quantitative PCR of mitochondrial to nuclear DNA expression (mtDNA/nDNA) in LV and RV myocytes from n = 3 animals. **G:** Pooled densitometric analysis and **H:** representative Western blots of protein expression of Mfn1 in LV and RV myocytes from n = 6 animals. Data are mean ± SEM. ETC, electron transport chain; Mfn1, mitofusin 1.

To determine whether there was an enzymatic basis for the difference in maximal OCR in LV and RV myocytes, we spectrophotometrically measured the activities of each electron transport chain (ETC) complex. While there was no significant difference in the activities of complexes I, II, and IV, complex V activity was 1.4 ± 0.15-fold higher in the RV myocytes compared to LV (*p* = 0.048, **Figure 2E**). Notably this was not due to significant differences in protein expression of ETC enzymes between myocytes of the RV and LV (**Figure 2C-D**).

Because mitochondrial numbers can account for differences in respiration, we next quantified mitochondrial (mt) DNA content by qPCR. We observed a trend toward greater mtDNA levels in RV myocytes compared to LV (*p* = 0.128, **Figure 2F**). Additionally, the levels of mitochondrial fusion protein Mitofusin1 was also found to be 1.33 ± 0.10-fold higher in RV myocytes compared to LV (*p* = 0.037, **Figure 2G-H**). These data suggest differences in the numbers and organization of mitochondria in RV myocytes that potentially underlie their elevated respiration rates.

### 3.3 Fatty acid metabolism supports enhanced RV maximal OCR

Glycolytic rate, as measured by the extracellular acidification rate (ECAR) was higher in cardiomyocytes isolated from RV compared to LV (1.7 ± 0.25 vs 0.95 ± 0.09 pH/min/mg protein, *p* = 0.014, **Figure 3A**). Pyruvate dehydrogenase activity did not significantly differ between RV cardiomyocytes compared to the LV (*p =* 0.22; **Figure 3B**).

**Figure 3.**
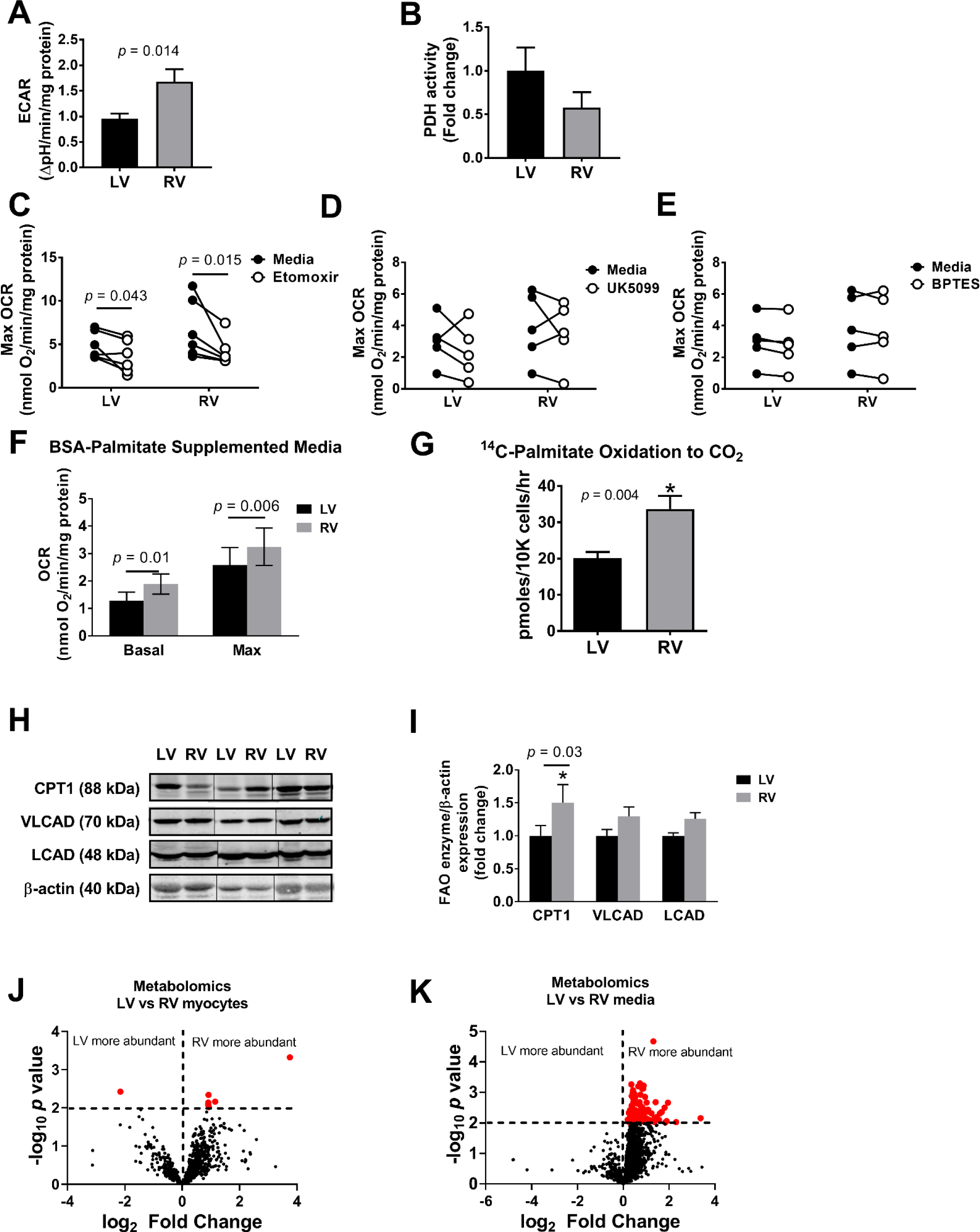
Distinct fuel utilization in LV and RV cardiomyocytes. **A:** Glycolytic rate, as measured by the extracellular acidification rate (ECAR) by extracellular flux analysis in LV and RV myocytes from n = 14 animals. **B:** Enzymatic activity of pyruvate dehydrogenase (PDH) in LV and RV myocytes from n = 5 animals. **C-E:** Quantification of the maximal OCR following injections of media or etomoxir, inhibitor of fatty acid oxidation; UK5099, inhibitor of glucose oxidation; and BPTES, inhibitor of glutaminolysis. **F:** Basal and maximal OCR in LV and RV cardiomyocytes from n = 10 animals supplemented with BSA-palmitate. **G:** Rate of ^14^C-palmitate oxidation to CO2 in LV and RV myocytes from n = 8 animals. **H:** Representative Western blots and **I:** pooled densitometric analysis of protein expression of carnitine palmitoyl transferase 1 (CPT1), very long chain acyl dehydrogenase (VLCAD), and long chain acyl dehydrogenase (LCAD) in LV and RV myocytes from n = 6 animals. **J-K:** Volcano plot visualization of 1,621 putative metabolites detected in LV and RV myocytes from n = 8 animals and media using untargeted LC-MS/MS analysis following normalization, practical filtration for missing values and applying Student’s two-tailed t-test. Data points in red represent significance of *p*-value < 0.01. Data are mean ± SEM.

To determine whether the increased maximal respiratory capacity in RV cardiomyocytes was due to a difference in the type of substrate being metabolized, we next measured maximal OCR in RV versus LV cardiomyocytes in the presence of inhibitors of pyruvate oxidation (UK5099; **Figure 3D**), glutaminolysis (BPTES; **Figure 3E**) or mitochondrial fatty acid entry (Etomoxir; **Figure 3C**). Consistent with the cardiac reliance on fatty acid oxidation, only etomoxir significantly inhibited OCR in cardiomyocytes and to a greater degree in the RV (*p* = 0.043 in LV, *p* = 0.0500 in RV; **Figure 3C**). Basal and maximal OCR measured in media supplemented with BSA-palmitate were both significantly higher in RV myocytes compared to LV (*p* = 0.01 for basal OCR, *p* = 0.006 for max OCR; **Figure 3F)**. Direct measurement of radiolabeled palmitate oxidation by the RV versus LV cardiomyocytes confirmed that the RV demonstrated greater rates of FAO than the LV cardiomyocytes (*p* = 0.004; **Figure 3G**). Compared to LV, there was increased RV expression of key FAO enzymes CPT1 (1.5 ± 0.28-fold, *p* = 0.027), VLCAD (1.29 ± 0.14-fold, *p* = 0.18), and LCAD (1.26 ± 0.09-fold, *p* = 0.24, **Figures 3H-I**). Taken together, these data suggest that RV myocytes exhibit enhanced fatty acid metabolism which support their enhanced maximal oxidative capacity.

LV and RV myocytes and the media from which they were suspended were subjected to metabolomic analysis by high resolution LC-MS. This untargeted analysis of 1,621 putative metabolites showed few metabolites which were significantly more abundant within RV myocytes compared to LV (**Figure 3J**). However, a large number of putative metabolites were found to be significantly more abundant in RV media compared to LV media (**Figure 3K**), further supporting fuel utilization differences between myocytes from the LV versus RV. We postulate that differences in metabolic rate or compensatory pathways may underlie this observation. This untargeted analysis requires further exploration to identify putative metabolites and specific metabolic pathways that differ between LV and RV myocytes.

## 4. DISCUSSION

Prior efforts to investigate the distinction between the RV and LV response to stress implicate differing metabolic regulation in the two ventricles as a potential basis for the divergent clinical course of RV versus LV failure.^8,26-28,34,35^ To our knowledge, a direct comparison of cellular metabolism between cardiomyocytes from the RV and LV has not previously been performed. Here, we adapt a Langendorff-free method^30^ to segmentally isolate primary adult rat cardiomyocytes from separated LV and RV, permitting comparison of bioenergetic profiles from the two ventricles. Our method yielded intact myocytes from both ventricles which exhibit distinct patterns of oxygen consumption wherein RV myocytes show enhanced respiratory reserve capacity compared to LV myocytes accompanied by differences in metabolic substrate utilization.

Previous work has shown that the LV and RV match O_2_ supply to demand via distinct mechanisms,^34,36,37^ illustrating the interventricular contrast in myocardial metabolism. Along with its lower resting workload and O_2_ requirement, the RV has lower resting coronary blood flow and O_2_ extraction rate than the LV.^27,36,38,39^ While both ventricles augment coronary blood flow to meet higher O_2_ demands, the RV can additionally recruit an O_2_ extraction reserve in response to increased work,^26^ inotropic stimulation,^27^ and exercise.^39^ Our study’s key finding that control RV myocytes exhibit higher FCCP-stimulated maximal oxygen consumption rate compared to LV myocytes, proposes a mitochondrial basis for the RV’s myocardial O_2_ extraction reserve.

In our study, RV myocyte spare respiratory capacity appeared to be dependent on FAO where RV myocyte maximal OCR was attenuated when endogenous FAO was inhibited with etomoxir. Further indicating their fatty acid-avidity, RV myocytes showed higher FAO enzyme expression and ^14^C_-_palmitate oxidation rate, and addition of exogenous palmitate increased maximal OCR in RV myocytes but not in LV. Thus, RV myocytes appear to utilize fatty acids to modulate respiration more efficiently than LV myocytes.

Mitochondrial respiration is regulated at multiple levels, including modulation of ETC enzyme kinetics and composition, intrinsic efficiency and integrity of mitochondria, organization of the mitochondrial network, and the cellular and mitochondrial microenvironment (for example, substrate availability).^40-42^ Consistent with prior studies that compared LV and RV mitochondrial enzyme composition,^35^ our examination of ETC enzyme activity and expression did not demonstrate ventricular differences in complex IV, the site of O_2_ reduction, nor complex I, the entry point of electron flux. Thus, the increased RV myocyte respiratory reserve is likely not attributable to baseline differences in ETC activity. OXPHOS can be modulated by mitochondrial dynamics, wherein mitochondrial fusion is associated with increased cellular respiration and coupling efficiency.^43-46^ In bovine RV compared to LV, Bruns et al. reported higher mitochondrial content by DNA copy number and morphometric analysis, as well as increased expression of pro-fusion Mfn1 mRNA, implicating enhanced mitochondrial fusion in the RV.^7,8^ Our finding of increased Mfn1 protein expression and increased mtDNA levels in RV compared to LV myocytes similarly support increased mitochondrial fusion and content in RV myocytes, which can potentially underlie their enhanced maximal respiratory capacity. Although we did not directly assess cardiomyocyte ATP production, we observed increased complex V enzyme activity in RV myocytes, in line with the enhanced energetics associated with mitochondrial fusion.^43-46^ Further investigation of mitochondrial morphology and dynamics in LV and RV cardiomyocytes will be informative.

In summary, we developed a method to isolate primary cardiomyocytes from separated LV and RV of adult rats to facilitate comparative assessment of mitochondrial metabolism between intact cells from the two ventricles. Our studies reveal inherent differences in cardiomyocyte metabolism between the LV and RV at rest. Future studies are needed to determine whether these baseline LV versus RV differences could translate to distinct patterns of mitochondrial adaptation to stress. Primary cardiomyocytes obtained by this procedure show promise as a biological system for study of mitochondrial metabolism in heart disease.

